# Unexpected properties of nucleotide condensates could facilitate the RNA world

**DOI:** 10.64898/2026.11.18.767475

**Authors:** Samuel Hauf, Lipika Baidya, Ryo Nakamura, Jacob L. Fine, Hung T. Nguyen, Alan M. Moses, Mirco Dindo

## Abstract

A key question in the RNA world hypothesis for the origin of life is the nature of the compartment that separates and protects the RNA from its environment. While membranes are a leading candidate for the initial compartment, biomolecular condensates have been proposed as an alternative, possibly simpler route to compartments. Here we show that *bona fide* liquid condensates form spontaneously when nucleotides (such as ATP) are mixed with divalent cations and a short-chain alcohol or diol in water. These condensates do not require any polymer to form, but can concentrate polymers like RNA and DNA, which show unexpected chemical behavior inside the condensates. Most surprisingly, RNA survives for days inside nucleotide condensates under acidic conditions that cause rapid and extensive hydrolysis in bulk. This effect is also observed for a phosphate-ester model substrate and therefore likely reflects the suppression of backbone cleavage. Coarse-grained simulations suggest that the condensate interior is bound together by magnesium-phosphate bridging interactions and is therefore highly depleted of water. At neutral pH, the same compartment reverses the course of a hammerhead ribozyme reaction, shifting it from cleavage toward religation. Finally, we observe that nucleotide droplets also spontaneously organize phospholipids at their surface, suggesting a possible evolutionary route from biomolecular condensates toward membranes. Thus, we believe that nucleotide condensates, which are built from the same building blocks as the genetic polymers themselves, overcome long-standing challenges with the RNA world hypothesis for the origin of life by protecting nucleic acids and reshaping their chemistry.

## Introduction

Nucleic acids are central to life and, most likely, to its very beginning. RNA in particular can encode information and catalyze the formation of peptide bonds [1], which is why it is frequently placed at the origin of life with the RNA world, a proposed stage at which RNA performed most functions important for life [2, 3]. A key challenge to the RNA world hypothesis is the need for compartmentalization [4]. Indeed, membranes are often thought to be a defining characteristic of all cells, including the first, although the nature of the first membranes is debated [5, 6, 7, 8, 9, 10]. Membranes create a distinct environment that encapsulates and protects biological molecules and allows biochemical reactions. How RNA in the RNA-world scenario could have assembled membranes, or whether membranes came first and allowed RNA to emerge as an information carrier, remains an open question [11, 12, 13].

Membraneless organelles formed by liquid-liquid phase separation (LLPS) have also been considered as compartments for the origin of life [14, 15, 16, 17]. Indeed, these so-called biomolecular condensates concentrate and protect RNA, especially during cellular distress [18, 19, 20, 21, 22], organizing the interior of living cells and compartmentalizing biochemistry [18, 23]. Recently, other important aspects of biomolecular condensates have received attention as well. Such condensates sustain enzymatic activity within their crowded interior, modify reaction rates, and can even move and reorganize as active droplets [24, 25, 26, 27, 28, 29, 30, 31, 32], all properties that make them attractive candidates for primordial compartments.

Condensates have been widely explored as routes to prebiotic protocells, with coacervates of peptides, nucleotides, and their mixtures shown to concentrate cargo and host early chemistry [33, 34, 35, 36, 37]. Condensates can also maintain distinct internal microenvironments that differ from the surrounding bulk solution [38, 39, 40]. Building on these properties, extensive efforts have established biomolecular condensates as models for prebiotic protocells: they can form from polymers such as polypeptides, polynucleotides and their mixtures [41, 42, 43]; they concentrate prebiotically relevant species such as magnesium and RNA; and they accelerate a range of reactions, from ribozyme catalysis to non-enzymatic RNA chemistry and even hydrolytic reactions [44, 45, 46, 47]. Recent theoretical work has proposed RNA condensates capable of polymerization as a route to both replication and compartmentalization [48]. Most recently, simplified condensate systems have been demonstrated experimentally. Short oligonucleotides with only a few peptide residues still coacervate [43, 49] and nucleotides themselves condense in the presence of polyethylene glycol (PEG) as a crowding agent [50, 51]. RNA within polymeric coacervates has also been shown to resist hydrolysis during wet-dry cycling [52].

Together, these advances indicate that condensates could have been pivotal at the origin of life. However, challenges to the demonstration of condensates as the primordial compartment remain. All of the systems above require a polymer, either as the phase-separating component or as the crowder that drives condensation, and the prebiotic origin of such polymers is itself uncertain. Further, the very backbone that makes nucleic acids functional also renders them fragile [53]. The phosphodiester bonds of RNA and DNA can be cleaved by water in acidic conditions, with abundant magnesium and elevated temperatures, sometimes invoked as drivers of polymerization at the origin of life, accelerating this cleavage [54, 55, 56]. This is a challenge to the RNA world hypothesis because abiotic RNA synthesis would likely be of limited efficiency, while hydrolysis would continuously degrade any polymers formed. A mechanism to protect nucleic acids from cleavage, regardless of the chemistry that assembled them, seems critical for life to emerge [57, 58, 59].

Here, we consider the simplest possible condensates formed from a single nucleotide (ATP) and show that they offer solutions to some of the challenges with the compartmentalization model. In water and in the presence of divalent cations and co-solvents such as short-chain alcohols (diols) of only a few carbon atoms (components plausibly available on the early Earth) [60, 61, 62, 53], we find that nucleotides form condensates in the absence of any polymer. We show that once formed, ATP condensates can recruit different kinds of RNA and DNA, creating a microenvironment that modulates nucleic acid behavior. In particular, a hammerhead ribozyme is shifted from cleavage toward religation, an effect previously reported in peptide-RNA coacervates [45]. Remarkably, in ATP condensates, RNA is protected from hydrolysis for days under acidic conditions that often cause rapid and extensive hydrolysis in bulk, overcoming a key challenge to the RNA world. Using coarse-grained simulations, we show that a protection mechanism arises from the strong depletion of water inside the condensates. Finally, we observe self-organization of a phospholipid layer around the ATP condensates, suggesting that nucleotide compartments may have the capacity to spontaneously develop membranes.

## Results

### Nucleotide condensates concentrate nucleic acids, whereas inorganic phosphate does not

Having observed that RNA undergoes phase separation in the presence of short-chain diols [63], we asked whether nucleotides themselves could be condensed in the same way, without any polymer. Mixing ATP with magnesium ions and short-chain diols produced micrometer-sized liquid droplets (see Methods). To test the relevance of nucleotide condensates for the RNA world, we first examined whether they had the capacity to recruit RNA. Indeed, ATP droplets concentrated single-stranded RNA and DNA, but also methylene blue, resorufin, and an ATP tracker, drawing each into the dense phase (Fig. 1a). This confirms that a nucleotide condensate is not merely a dense phase, but a separate environment that other organic molecules can enter.

**Figure 1:**
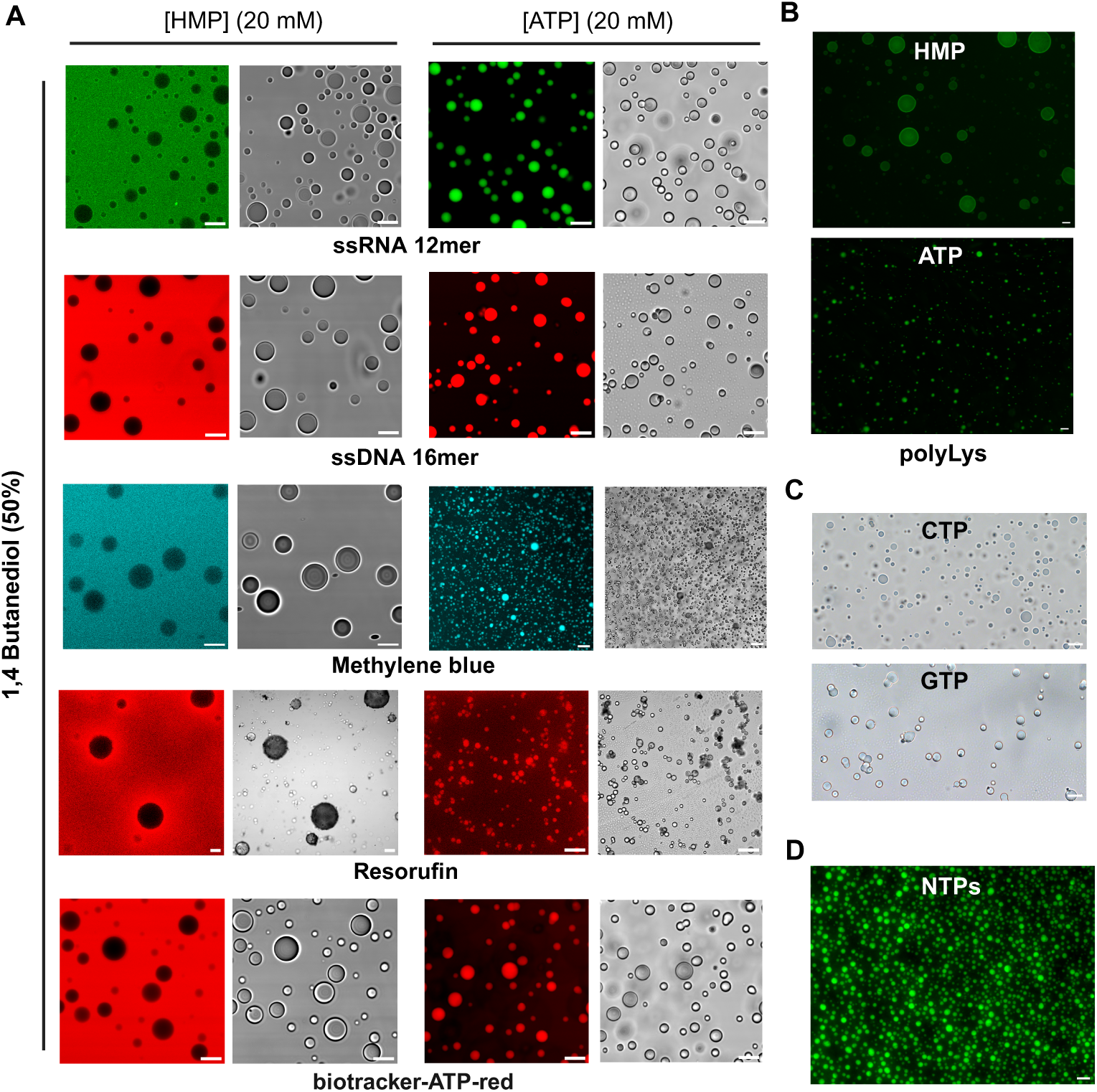
Nucleotide condensates concentrate nucleic acids and small molecules, whereas inorganic phosphate condensates do not. (**a**) Partitioning of five cargoes into HMP versus ATP droplets (20 mM HMP or 20 mM ATP, 50% 1,4-butanediol, 30 min): FAM-labeled ssRNA 12-mer (green), Cy3-labeled ssDNA 16-mer (red), methylene blue, resorufin and a biotracker ATP-red tracker are all excluded from HMP (left pair of panels) and concentrated in ATP (right pair of panels). Pairs of panels correspond to fluorescence and corresponding brightfield. **(b)** Exclusion from the HMP phase is not simple impermeability. Positively charged poly-L-lysine partitions into both HMP (top panel) and ATP (bottom panel) droplets (obtained with 1,4-butanediol and ethanol, respectively) entering even the otherwise exclusionary phosphate phase. **(c)** Condensation extends to other nucleoside triphosphates: CTP and GTP both form droplets (GTP after acidification), shown here in brightfield. **(d)** Condensation is also a property of nucleotide mixtures. An equimolar NTP pool (10 mM each nucleotide, 40 mM total; 20 mM MgCl_2_, 50% 1,4-butanediol) forms droplets and takes up a FAM-labeled ssRNA 16-mer (green). Scale bars, 10 or 20 µm unless otherwise indicated; acidic conditions throughout unless noted (pH *<* 3).

Nucleotide condensation has so far been reported only for ATP [50, 51]. However, in a prebiotic setting, a mixture of nucleotides rather than a single purified species may have been present. Therefore, we tested other nucleotides and found that CTP and GTP also formed droplets (Fig. 1c). Phase separation was also possible using equimolar mixtures of the four natural NTPs, and these condensates also took up RNA (Fig. 1d). Condensation is therefore a general feature of nucleoside triphosphates rather than of ATP alone.

To determine whether the enrichment of RNA and DNA inside the condensates was a specific property of nucleotide condensates or a property of other components of the system, we tested phase-separated inorganic hexametaphosphate (HMP) under the same conditions. HMP is a simple polymeric phosphate that can form abiotically by heating orthophosphate and may have formed on the early Earth through meteorite impacts or volcanism acting on phosphate-bearing minerals [64, 65, 66]. We observed that HMP demixed even more readily than nucleotides. Indeed, HMP droplets formed with every diol tested under acidic, neutral and basic conditions (Fig. S1), although ethylene glycol required a higher HMP concentration (see Methods).

We next tested whether phosphate, rather than the nucleobase, drives the phase separation. Neither adenine nor adenosine condensed under any condition tested, whereas sodium triphosphate, another inorganic phosphate, condensed readily (Fig. S2). This is consistent with a large body of work on phase separation of inorganic phosphates [67, 68, 69, 70, 66].

However, HMP droplets, identified in brightfield microscopy, excluded most solutes: RNA, DNA and small organic molecules were largely absent from their interior in the fluorescence channels (Fig. 1a). Thus, phase separation per se is not specific to nucleotides. Under conditions where both phosphate species (ATP and HMP) form a dense liquid phase, only the nucleotide readily recruits other molecules. The exclusion from the HMP phase was not simply due to impermeability: positively charged poly-L-lysine partitioned into both condensates (HMP and ATP; Fig. 1b), consistent with a negatively charged, phosphate-rich interior that can form a complex coacervate with a polycation. A positive charge can therefore carry molecules into the phosphate phase. Thus, in this system, phase separation requires the phosphate, whereas a compartment capable of holding RNA or other organic molecules appears to additionally depend on the nucleoside moiety. We propose that demixing and cargo uptake are separable properties, each governed by a different part of the molecule.

### Diols facilitating nucleotide condensation avoid adjacent hydroxyl groups

Nucleotide phase separation has previously been reported under two different regimes: at molar concentrations of both nucleotide and divalent cation (around 0.7-1 M) [71], where ATP and Mg^2+^ assemble into a percolated, gel-like phase, and at lower nucleotide concentrations in the presence of a polymer crowder such as PEG [50, 51]. In both cases a polymer or a near-molar nucleotide reservoir is required to drive the transition. To our knowledge, the formation of liquid nucleotide condensates at millimolar nucleotide and magnesium concentrations, driven by a short-chain diol or alcohol in the absence of any polymer, has not been described before. Alcohols and diols differ widely in chain length and hydroxyl positioning, identifying which of them support nucleotide condensation, and why, can reveal what drives the phase separation process. We therefore compared a series of chemically related co-solvents. Diols and alcohols act here as co-solvents rather than macromolecular crowders. Their effect depends on the position of the hydroxyl groups rather than on molecular size. Indeed, 1,2– and 1,3-propanediol have similar sizes but give aggregates and droplets, respectively, which argues against a simple excluded-volume mechanism. We propose that short alcohols lower the water activity and dielectric constant of the medium, favoring phosphate-Mg2+ bridging, whereas PEG crowds the medium by excluding volume. Diols were not included in our simulations, so this mechanism remains to be tested directly. Whereas inorganic phosphate condensed with every diol or alcohol tested, ATP proved far more selective, and the pattern of co-solvents that triggered its phase separation was itself informative (Fig. 2a,b). Both methanol and ethanol supported ATP condensation, indicating that between these two simple alcohols (a single hydroxyl group), chain length was immaterial. Among diols, by contrast, the position of the hydroxyl groups determined the outcome. Vicinal diols, with hydroxyls on adjacent carbons, either failed to trigger condensation (ethylene glycol, glycerol) or produced aggregates (1,2-propanediol). Surprisingly, separating the hydroxyls by even one carbon restored droplet formation (1,3-propanediol) (Fig. 2b). The length of the carbon chain between non-vicinal hydroxyls did not further affect the outcome across the diols tested (1,4-butanediol, 1,6-hexanediol). This selectivity points to requirements of nucleotide phase separation beyond desolvation. A short alcohol or diol lowers the water activity and dielectric constant of the medium, weakening the hydration of the charged groups and favoring association. For the phosphate groups alone, these effects are sufficient to trigger phase separation. In the case of the nucleotide, its base and its magnesium-bridged phosphates must be brought into register, and vicinal hydroxyls, which hydrate cooperatively and can chelate divalent cations, probably interfere with this process (Fig. 3a).

**Figure 2:**
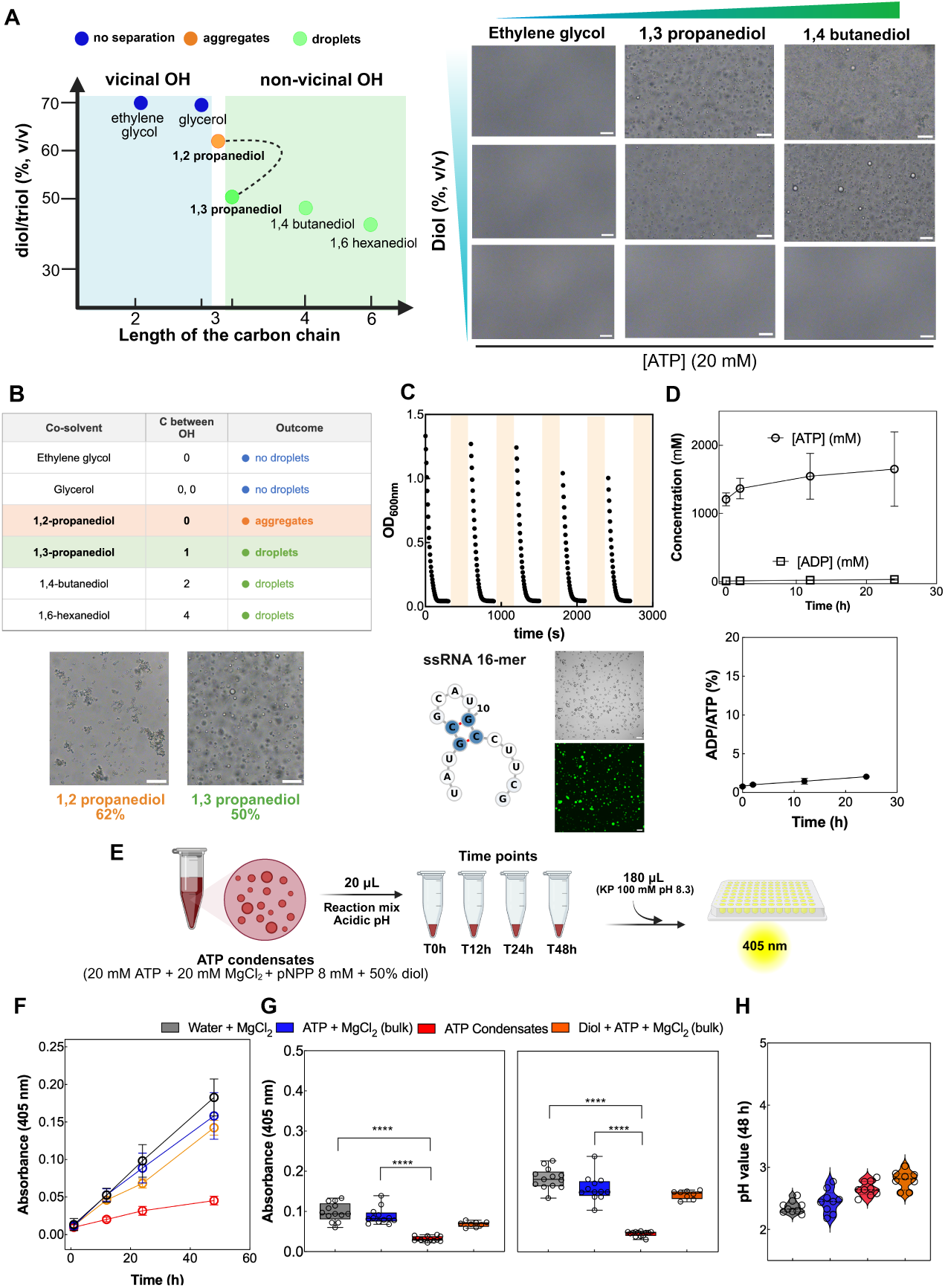
Nucleotide condensation follows a structural rule, is reversible, and suppresses a model phosphoester hydrolysis. (**a**) Co-solvent structure–activity map. The spacing of the hydroxyl groups, not chain length, decides the outcome (no separation, aggregates, or droplets). Representative brightfield images of ATP condensation at 20 mM ATP. **(b)** Co-solvents grouped by carbons between the hydroxyls, with representative images of the propanediol isomers (1,2-propanediol, 62%, aggregates; 1,3-propanediol, 50%, droplets). **(c)** Reversibility. Cooling (∼1 *^◦^*C) induces condensation and RNA uptake, and warming (30 *^◦^*C) dissolves the droplets, reversibly over five cycles. Brightfield and confocal images show the ssRNA 16-mer (∼5 µM) inside cold-formed condensates. **(d)** ATP and ADP in ATP condensates quantified by HPLC at 0, 2, 12 and 24 h (top). ADP/ATP ratio stays at or below 2% (bottom). Mean ± SD of two biological replicates with a technical measurement at a different dilution (*n* = 3 for 0, 2 and 12 h; *n* = 2 for 24 h). **(e)** Model phosphoester hydrolysis assay. *Para*-nitrophenyl phosphate (pNPP, 8 mM) was incubated in ATP condensates; aliquots were developed in potassium phosphate and *p*-nitrophenol was read at 405 nm. **(f)** Absorbance at 405 nm over 48 h for water + MgCl_2_, ATP + MgCl_2_ bulk, ATP condensates and diol + ATP + MgCl_2_ bulk. Release is suppressed only in condensates (*p <* 0.0001 in all comparisons; Welch’s one-way ANOVA with Dunnett’s T3). *n* = 13, 12, 13 and 8, pooled across independent experiments. **(g)** Absorbance at 405 nm at 24 h (left) and 48 h (right) for the same conditions (∗ ∗ ∗∗, *p <* 0.0001; same test). **(h)** pH at 48 h for the four conditions (*n* = 3 each). All remain comparably acidic, so the suppression is not due to pH. Scale^6^bars, 20 µm unless otherwise indicated.

**Figure 3:**
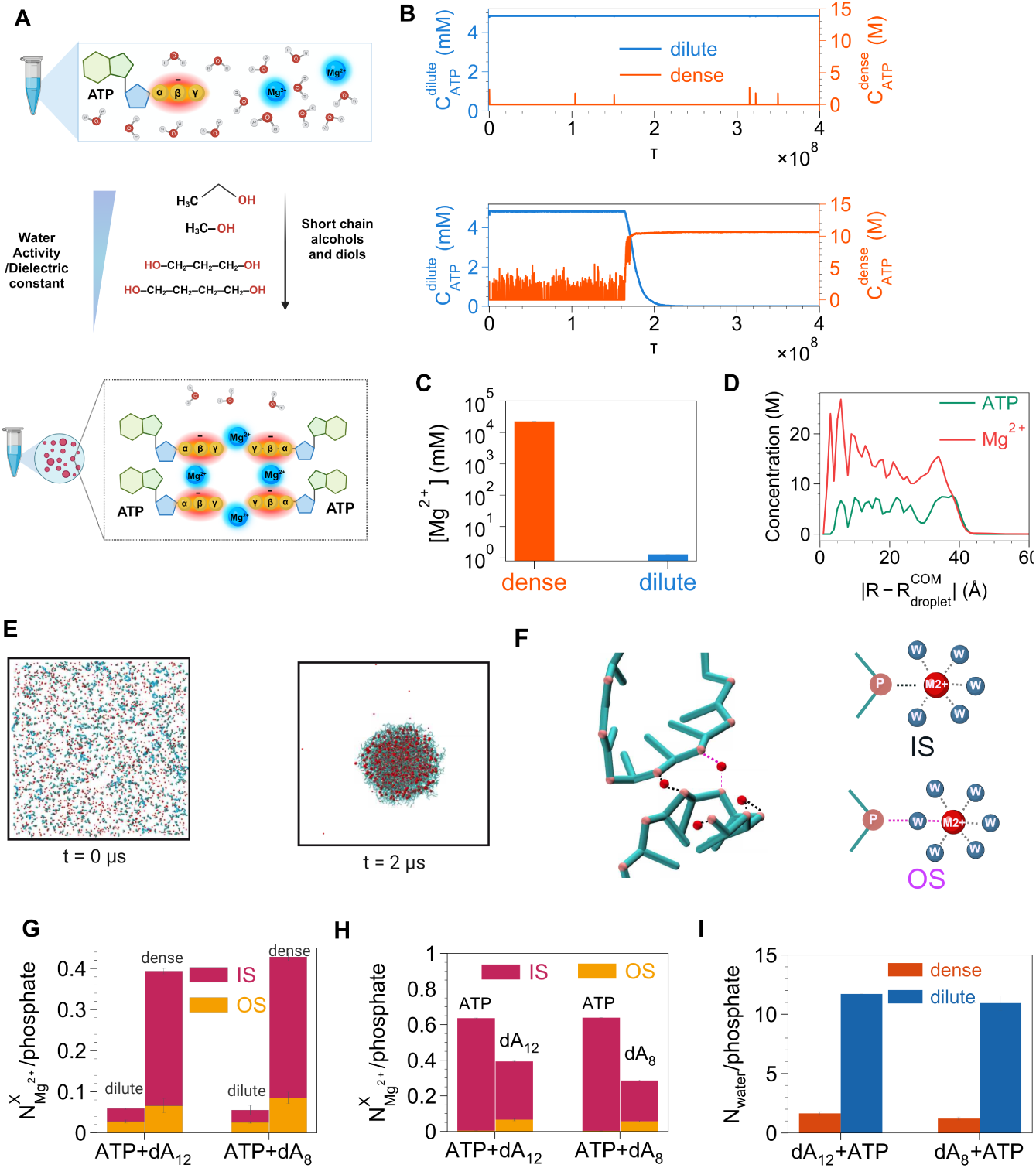
Simulations suggest a magnesium-bridged interior with reduced water availability. (**a**) Conceptual scheme: a short-chain alcohol or diol lowers the water activity and dielectric constant of the medium, favoring association of ATP and Mg^2+^, while vicinal hydroxyls interfere with base stacking and cation bridging. **(b)** Coarse-grained molecular simulations of ATP. Without Mg^2+^ (top) ATP stays in the dilute phase and does not condense; with 10 mM Mg^2+^ (bottom) ATP partitions into a dense phase over time (initial ATP 5 mM), with representative snapshots (ATP, cyan; Mg^2+^, red). **(c)** Mg^2+^ is enriched within the condensate relative to the surrounding solution (initial Mg^2+^ 10 mM). **(d)** Radial concentration of ATP phosphate and Mg^2+^ from the condensate center of mass, showing the two colocalize across the dense phase, consistent with Mg^2+^–phosphate bridging. **(e)** Simulation of ATP with ssDNA (dA_12_) and Mg^2+^ (initial ATP 5 mM, DNA 300 µM, Mg^2+^ 10 mM): snapshots at *t* = 0 and 2 µs show co-condensation.**(f)** Close-up and schematic of Mg^2+^ coordination to a phosphate: inner-shell (IS), direct coordination; outer-shell (OS), coordination through a bridging water. **(g)** IS versus OS Mg^2+^ per phosphate in the dilute and dense phases (ATP+dA_12_, ATP+dA_8_): within the condensate more than 80% of bound Mg^2+^ is inner-shell (dehydrated). **(h)** Mg^2+^ bound per phosphate for ATP versus DNA (dA_12_, dA_8_): Mg^2+^ associates roughly twofold more with ATP than with the DNA backbone. **(i)** Number of hydration water per phosphate in the dilute and dense phases (ATP+dA_12_, ATP+dA_8_).

### Chemical properties of ATP condensates

A protocell-like compartment must be able to form and dissolve repeatedly, for instance under temperature cycles, so that its contents can mix and be redistributed, and it should work at near neutral pH, where RNA catalysis is possible. We therefore first tested whether condensation was reversible and responsive to temperature. At near-neutral pH, using a condition below the threshold for phase separation at room temperature (32 mM ATP, 80 mM MgCl_2_, 125 mM Tris-HCl, pH 8, 100 mM NaCl, 30–35% 1,4-butanediol), no droplets formed, whereas cooling to near 1 *^◦^*C on ice induced condensate formation. Under near-neutral conditions, MgCl_2_ (80 mM) is in large excess over ATP (32 mM) and RNA (5 µM), so only a fraction of the MgCl_2_ is expected to be bound to either ATP or RNA. These cold-formed condensates concentrated RNA as did the condensates formed under acidic conditions, showing that the ability to take up RNA is retained at low temperature and near-neutral pH (Fig. 2c). While warming to 30 *^◦^*C the droplets dissolved, whereupon the cycle could be repeated for at least four more rounds (Fig. 2c). Condensation is therefore a genuine, reversible phase transition rather than irreversible precipitation, consistent with the upper critical solution behavior reported for nucleotide condensates [50] and here extended to near-freezing temperatures.

Next, we explored the chemical properties of the ATP condensates. Under our standard condition (20 mM ATP, 20 mM MgCl_2_, 50% 1,4-butanediol), the nucleotide was strongly enriched in the dense phase. HPLC quantification gave an ATP concentration of 1.2–1.7 M in the dense phase against ∼11 mM in the supernatant (Fig. 2d), a ∼100-fold enrichment (dense-phase volume ∼5–8 *µ*L per mL of reaction, measured directly by pipetting after centrifugation).

We also measured ATP and ADP within the dense phase to test whether the concentrated nucleotide was being hydrolyzed. ATP in the dense phase remained in the molar range at all time points measured (0, 2, 12 and 24 h). ADP stayed in the low millimolar range (9–34 mM), and the ADP/ATP ratio remained at or below 2–3% throughout, indicating that ATP within the condensate is not appreciably hydrolyzed over 24 h (Fig. 2d).

We next asked whether the stability we observed for the nucleotide itself extended to other phosphate esters. Para-nitrophenyl phosphate (pNPP, 8 mM) was incubated in ATP condensates (20 mM ATP, 20 mM MgCl_2_, 45–50% 1,4-butanediol, acidic pH) and its hydrolysis followed spectrophotometrically as the release of p-nitrophenol at 405 nm (Fig. 2e). Over 48 h, hydrolysis was markedly suppressed in the condensates relative to all bulk conditions tested (water + MgCl_2_; ATP + MgCl_2_; ATP + MgCl_2_ with diol below the condensation threshold; Fig. 2f, g; *p <* 0.0001 at both 24 and 48 h, Welch’s one-way ANOVA with Dunnett’s T3). The pH remained comparably acidic across all conditions (Fig. 2h), ruling out a pH artifact. Because no protein is present in this assay, the suppression reflects an intrinsic chemical property of the dense phase that slows phosphate-ester hydrolysis.

### Simulations suggest a magnesium-bridged interior with reduced water availability

To resolve the physicochemical basis of nucleotide condensation (Fig. 3a), we turned to coarse-grained molecular simulations. First, we investigated the role of phosphates in driving phase separation by performing simulations of ATP (5 mM) in the absence and presence of Mg^2+^ ions. Without magnesium, ATP remained dispersed in the dilute phase and did not form any condensates (Fig. 3b, top). However, addition of 10 mM Mg^2+^ drove it into a dense phase over the course of the simulation (Fig. 3b, bottom). The dense-phase ATP concentration reached ≈10 M, higher than the 1.2–1.7 M measured experimentally (Fig. 2d), but consistent with strong enrichment of ATP in the dense phase. Magnesium was also strongly enriched within the dense phase relative to the surrounding solution (Fig. 3c). This observation suggests that ATP condensate formation is driven by phosphate-Mg^2+^ bridging. We confirmed this by tracking the radial distribution of ATP phosphates and Mg^2+^, which were co-localized throughout the condensate (Fig. 3d). In our simulations, cation bridging of the phosphates is therefore the structural basis of condensation, similar to what has been observed in RNA folding [72, 73, 74].

To characterize the microenvironment of the dense phase when nucleic acids are present we performed simulations with ssDNA (dA_8_ or dA_12_) added to the ATP–Mg^2+^ system. The DNA co-condensed with ATP and magnesium (Fig. 3e), consistent with our experiments showing that ATP condensates take up single-stranded RNA and DNA (Fig. 1a). Remarkably, we observed that magnesium bound to the nucleic-acid backbone significantly shifted its coordination (Fig. 3f). In the dilute phase, Mg^2+^ coordinated phosphate through both inner-shell (direct) and outer-shell (water-bridged) modes, whereas within the dense phase more than 80% of bound Mg^2+^ was inner-shell (Fig. 3g). The predominance of inner-shell, dehydrated magnesium hints at a condensate interior in which water is much less available. Because hydrolysis of phosphate esters and phosphoanhydrides depends on water to mediate proton transfer and stabilize the transition state (either as the nucleophile or, for RNA cleavage via the 2’-OH), such a water-depleted environment offers a plausible basis for suppression of hydrolysis.

To corroborate the change in water content within the condensate, we estimated the average number of water molecules associated with the phosphates in the dilute and dense phases from the simulated solvent-accessible surface area (SASA) (Fig. S3). We observed an approximately 7–9-fold reduction in the estimated water content within the condensate compared with the dilute phase (Fig. 3i). This result is consistent with a condensate interior that is substantially depleted of water, which could create a protective environment that reduces the susceptibility of nucleic acids to hydrolysis. In addition, magnesium associated roughly twice as strongly with ATP as with the DNA backbone (Fig. 3h), leaving comparatively less ions around the nucleic acid. Since Mg^2+^ can catalyze phosphodiester bond cleavage, the lower availability of magnesium around DNA may also contribute to its slower hydrolysis.

### Nucleic acids are protected from hydrolysis within the compartment

Nucleotide condensates thus have the following properties. They concentrate nucleic acids (Fig. 1), suppress the hydrolysis of a model phosphomonoester in the absence of any protein component (*para*-nitrophenyl phosphate; Fig. 2e–h), and in coarse-grained simulations they form a magnesium-bridged interior in which the cations are largely dehydrated, with far less water per phosphate in the dense phase (Fig. 3). The obvious next question was whether nucleic acids held inside such compartments would also be protected from cleavage.

We tested the stability of three short RNAs, a single-stranded 12-mer, a single-stranded 20-mer and a structured 28-mer hairpin. As expected, each was concentrated into the ATP condensate (Fig. 4a, d, confocal). Confocal imaging further revealed that structured and tailed RNAs do not fill the droplet uniformly but concentrate into discrete intradroplet domains, as also seen for a second RNA hairpin and for the poly-A-tailed hammerhead substrate (Fig. S4), indicating that nucleic-acid uptake proceeds alongside a sub-droplet organization that recalls multiphase condensate behavior.

**Figure 4:**
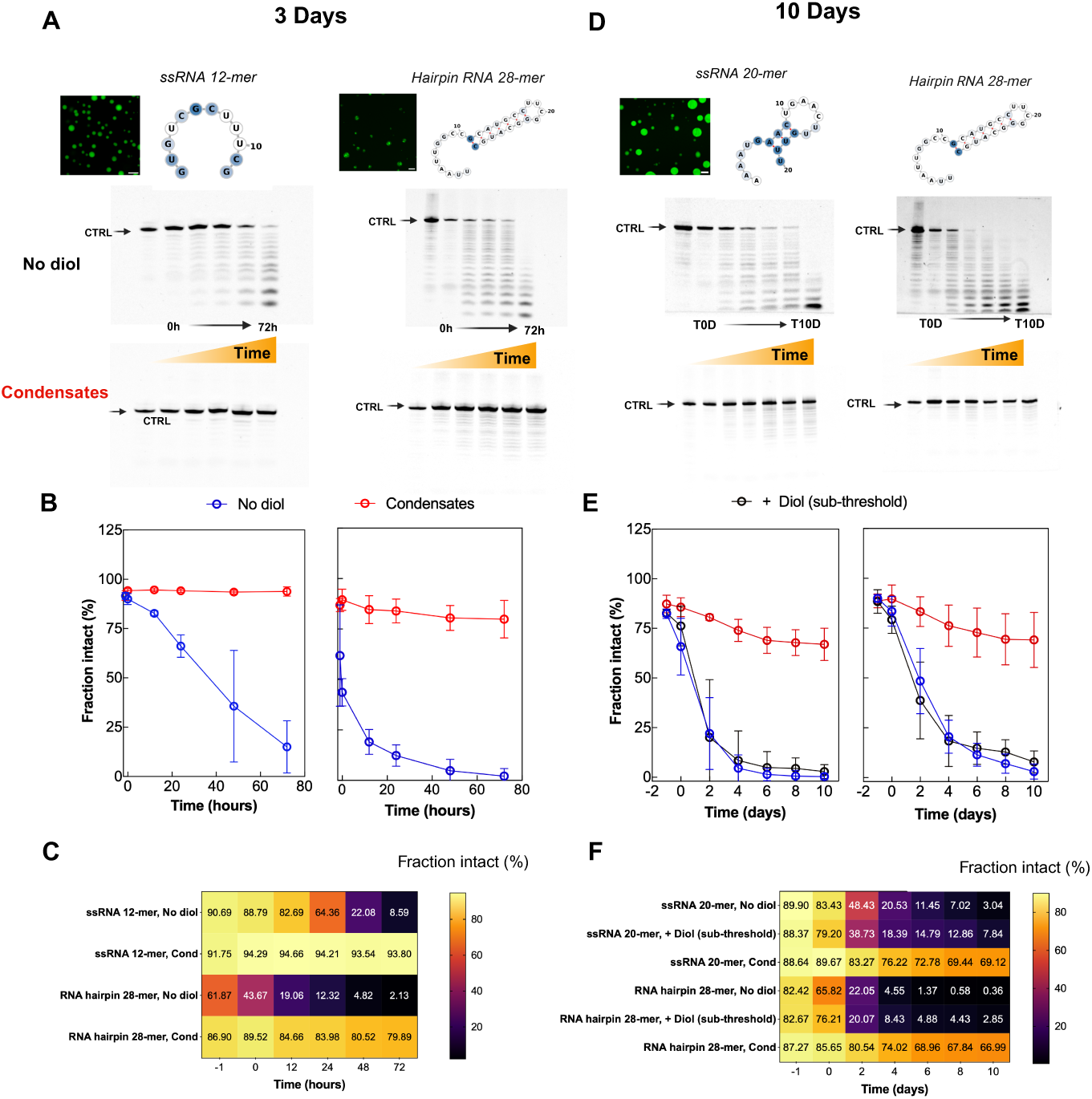
RNAs are protected from hydrolysis inside nucleotide condensates, over hours and over days. (**a**) Short time course (0–72 h) for the ssRNA 12-mer and the 28-mer hairpin. Top, confocal fluorescence images showing the RNA concentrated inside ATP condensates, together with the secondary-structure prediction (RNAfold, minimum free energy with the “avoid isolated base pairs” option; RNA Turner 2004 parameters, 1 M salt, 37 *^◦^*C [75, 76]). Bottom, negative images of the fluorescence signal after denaturing urea-PAGE for the no-diol control and the condensate, with untreated control (CTRL). **(b)** Fraction of intact oligonucleotide over 72 h for the ssRNA 12-mer (left) and the 28-mer hairpin (right), no-diol control (blue) versus condensate (red); mean ± SD, *n* = 4. **(c)** Heatmap of the fraction intact (%) from −1 to 72 h for the two oligonucleotides in (a). Inside the condensate the ssRNA 12-mer stays above 90%, and the 28-mer hairpin declines from about 87% to about 80%. In the no-diol control the ssRNA 12-mer falls to about 9% and the 28-mer hairpin to about 2% by 72 h. **(d)** Long time course (0–10 days at 37 *^◦^*C) for the ssRNA 20-mer and the 28-mer hairpin, with the same layout as in (a): confocal images and structure prediction (top), denaturing urea-PAGE (bottom). **(e)** Fraction of intact oligonucleotide over 10 days for the ssRNA 20-mer (left) and the 28-mer hairpin (right) in three conditions: condensate (red), no-diol control (blue), and water + ATP + MgCl_2_ with 1,4-butanediol below the condensation threshold (black); mean ± SD, *n* = 4. **(f)** Heatmap of the fraction intact (%) from −1 to 10 days for the two oligonucleotides and the three conditions in (e). At day ten the condensate keeps the ssRNA 20-mer at about 69% and the 28-mer hairpin at about 67%, whereas in the no-diol control the ssRNA 20-mer falls to about 3% and the hairpin to below 1%. Sub-threshold diol (no condensate formed) gives only partial protection. Throughout, “no diol” denotes a tube containing water, ATP and MgCl_2_ at the same concentrations as the condensate but without 1,4-butanediol, so no condensate forms; “+ diol (sub-threshold)” denotes a tube in which 1,4-butanediol is added at 35% (v/v), a concentration insufficient to drive condensation; “condensate” denotes the fully condensed system (50% 1,4-butanediol). RNAs at 5 *µ*M; 20 mM A1T0 P, 20 mM MgCl_2_; acidic conditions (pH 2–2.3). Scale bars, 20 *µ*m.

In the no-diol control (water + ATP + Mg^2+^, acidic pH, no condensate formed), both RNAs tested, the 12-mer ssRNA and the 28-mer hairpin, broke down steadily over 72 h, with most of the full-length material lost within that time on denaturing urea-PAGE gels. Remarkably, the same molecules remained largely intact over the same period inside the condensate phase (Fig. 4a). Over the course of the experiment, full-length material in the no-diol control dropped to about 9% for the 12-mer ssRNA and to about 2% for the hairpin by 72 h, while in the condensate the 12-mer stayed above 90% and the hairpin declined only from about 87% to about 80% (Fig. 4b, c).

That a small phosphomonoester (*para*-nitrophenyl phosphate; Fig. 2e–h) and RNAs of different length and fold are protected to much the same extent argues against simple steric shielding of a given structure. Further, the degradation pattern on the gel is a complete ladder with bands at every nucleotide position across the sequence (Fig. 4a, d), consistent with random non-enzymatic phosphodiester cleavage via the 2’-OH intramolecular mechanism. An RNase A contamination would instead give a partial ladder restricted to pyrimidine positions, which is not what we observe. Moreover, the suppression of *para*-nitrophenyl phosphate hydrolysis (Fig. 2e–h) occurs in a reaction that contains no protein at all. The same protection extends to ssDNA (Fig. S5), which lacks the 2’-OH and therefore degrades in acidic bulk through a different chemistry than RNA, yet is similarly stabilized inside the condensate. Together, these observations indicate that the stabilization reflects a chemical effect of the dense phase, not the exclusion of trace amounts of nucleases. We propose that the condensate slows cleavage of the phosphodiester backbone, and that this is an intrinsic property of the environment inside the condensate itself.

Having demonstrated the stabilizing effect of ATP condensates at relatively short times, we next extended the incubation from three to ten days. This left the picture essentially unchanged (Fig. 4d–f). In the no-diol control, the 20-mer ssRNA was reduced to about 3% of full-length material by day ten and the hairpin to under one percent. Adding diol at a concentration below the condensation threshold slowed this only partially (Fig. S6), confirming that the protection reflects the condensed phase environment and not an intrinsic chemical effect of the diol itself. In the condensates, the 20-mer and the hairpin fell from about 89% and 87% to about 69% and about 67% intact, respectively. The condensate samples show a modest decline over the same period, but without the shorter products seen in the no-diol control and in the sub-threshold diol sample, so this decrease may reflect technical factors rather than hydrolysis. Also, denaturing gels of the supernatant from the condensate samples showed no detectable RNA or degradation products, indicating that the fragments did not leave the droplet (data not shown). Because these experiments were run at 37 *^◦^*C in closed tubes, the compartment kept essentially the same volume throughout, unlike protection schemes that rely on drying, where the compartment itself is lost as the sample dehydrates [52]. This raised a further question of whether the condensates might do more than simply slow the hydrolysis of what they contain.

### Confinement reshapes catalysis

Given the stability of the condensates and the nucleic acids within them, we next tested the catalytic activity of a minimal hammerhead ribozyme inside the condensates. We incubated the ribozyme with its substrate at neutral pH, both in buffer and in the condensates, at two ribozyme concentrations (∼0.75 and ∼2.9 µM; Fig. 5a, b). In buffer, the cleaved fraction rose and then saturated, consistent with a reaction proceeding to completion. In contrast, in the condensates, the fraction of cleaved substrate reached a peak at around two hours and subsequently decreased, accompanied by the reappearance of uncleaved substrate. Substrate recovery was negligible in buffer, but reached around 45–53% inside condensates. Therefore, in the dense phase, the reaction did not simply run to completion as the equilibrium shifted to a more favored reverse direction, potentially due to the increased local concentration of the cleavage products within the condensate (Fig. 5c; see Discussion).

**Figure 5:**
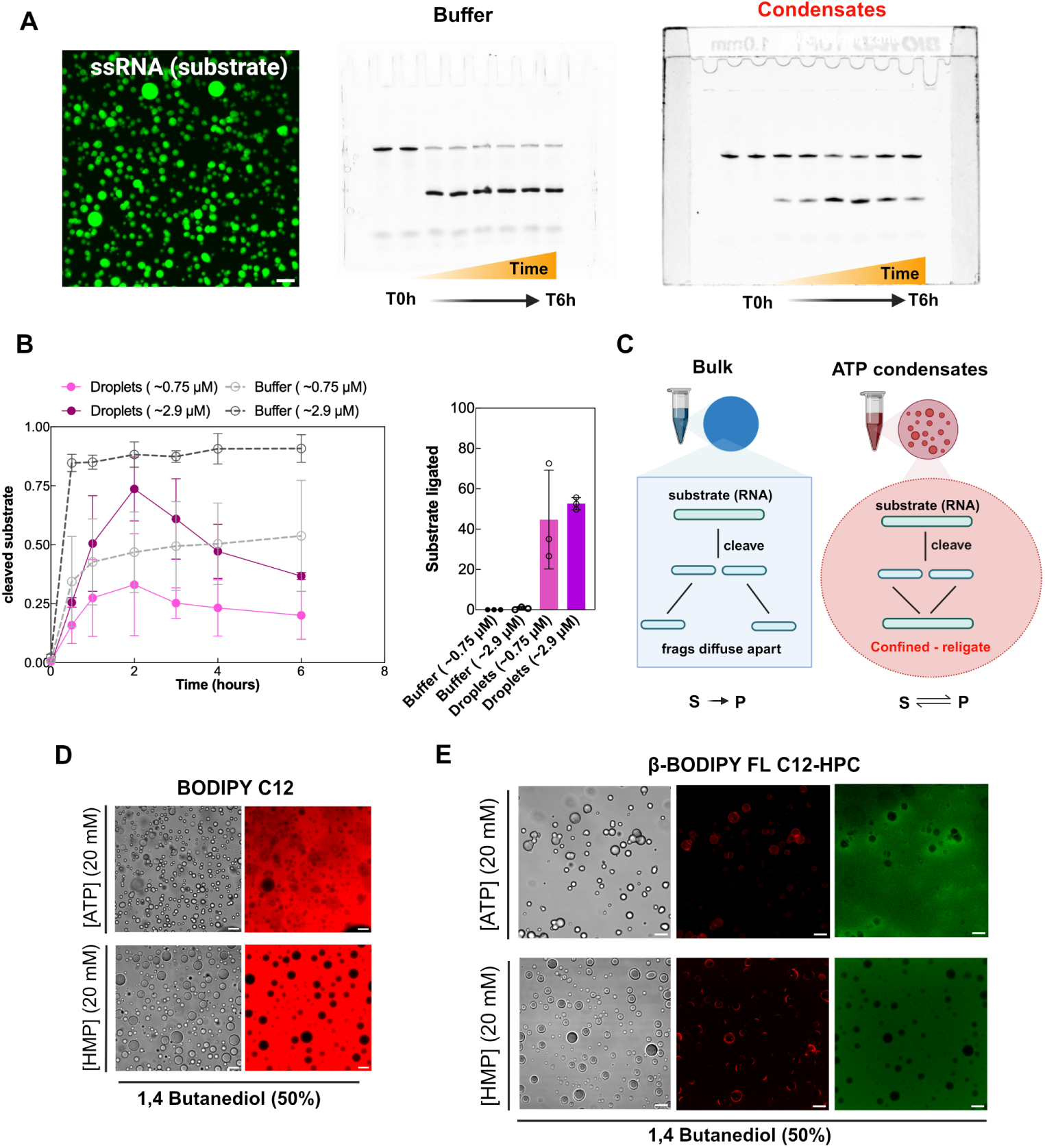
Confinement reshapes ribozyme catalysis, and the condensates spontaneously organize phospholipids at their surface. (**a**) Hammerhead ribozyme assay. Left, confocal image of the FAM-labeled substrate concentrated inside ATP condensates. Right, negative fluorescence images of denaturing gels (0–6 h) of the ribozyme reaction in buffer and in ATP condensates, at neutral pH (6.7–7.2), run at a high (∼2.9 µM) ribozyme concentration. Only the FAM-labeled substrate is visible on the gels. **(b)** Fraction cleaved over time, in buffer and in condensates at low (∼0.75 µM) and high (∼2.9 µM) ribozyme concentration. In buffer the cleaved fraction rises to a plateau, whereas in condensates it peaks near 2 h and declines as substrate is regenerated. Right, substrate recovery between 2 and 6 h, ∼45–53% in condensates and near zero in buffer. *n* = 3 independent experiments per condition; mean ± SD. **(c)** Scheme. In bulk, cleavage products diffuse apart and the reaction runs to completion (S→P); under confinement they remain proximal and religate, so the reaction becomes reversible (S⇌P). **(d)** The fatty-acid analogue BODIPY C_12_ is excluded from the interior of both ATP and HMP droplets (brightfield and fluorescence).**(e)** The phospholipid *β*-BODIPY FL C_12_-HPC accumulates at the interface of both ATP and HMP droplets (brightfield, red and green channels; the red signal decorates the interface while the green signal is distributed in the bulk).Scale bars, 20 µm.

Alongside their effect on catalysis, we also observed a remarkable phenomenon at the condensate boundary. The phospholipid *β*-BODIPY FL C_12_-HPC accumulated at the surface of both ATP and HMP droplets (Fig. 5e), while the fatty-acid analogue BODIPY C_12_ was excluded from the interior of both (Fig. 5d). The accumulation at the interface built up over hours, with the phospholipid initially distributed as foci and progressively spreading to a ring at the droplet surface by 24 h (Fig. S7). A compartment built from a single nucleotide can therefore not only concentrate, protect, and reshape nucleic-acid chemistry but also spontaneously organize phospholipids at its boundary, hinting at a possible route from a nucleotide compartment toward a membrane.

## Discussion

It is becoming clear that phase separation is a widespread phenomenon with potential significance for the origin of life. Although biological phase separation has been most widely appreciated in the context of polymers with weak multivalent interactions [77, 18], we find that nucleotides can form bona fide compartments, capable of sequestering organic molecules, organizing lipids, and protecting nucleic acids from degradation. This is in contrast to inorganic phosphate droplets, which are also liquid, coalesce on contact, yet exclude many biologically relevant molecules. We propose that the phase separation and partitioning properties of nucleotide compartments are the result of the structure of the nucleotide itself, which is composed of distinct moieties (phosphate, sugar, and base), each bestowing different chemical properties on different parts of the nucleotide molecule. The phosphate could drive liquid-liquid phase separation while the sugar and base could create a more hydrophobic environment capable of partitioning other organic molecules. Together, the combination of these different properties enables the formation of complex condensate compartments.

The effects the nucleotide compartment exerts on its contents (e.g. protection from hydrolysis and shifting of a reaction equilibrium) follow from the physical character of the dense phase by two distinct routes. The protection is consistent with reduced water activity inside the condensate [78], inferred from the predominantly dehydrated coordination of magnesium in our simulations (Fig. 3). Because hydrolysis of phosphoanhydride and phosphate-ester bonds depends on water, reduced water availability is expected to slow down backbone cleavage. The reversal of the ribozyme reaction, in contrast, is most likely due to confinement in condensates. Cleavage products that would otherwise diffuse apart in dilute solution remain in proximity in the dense phase and therefore easier to re-ligate. In this way, a transesterification reaction that normally runs to completion in bulk approaches an equilibrium in condensates.

The observed stabilization of RNA inside nucleotide condensates was surprising. Coacervates generally accelerate the reactions of the molecules they concentrate, by raising local reactant and catalyst concentrations, stabilizing intermediates, and providing molecular crowding [34, 35]. Hydrolytic reactions should be no exception, and ester hydrolysis in particular has been reported to accelerate inside coacervate microreactors [35]. A dense phase rich in phosphate and magnesium, both of which participate in the hydrolysis of phosphate esters in nucleic acids, might therefore be expected to accelerate their degradation. However, we observe the opposite. We previously found that RNA condensates neither markedly accelerated nor suppressed the hydrolysis of a nitrophenyl ester relative to bulk [63]; but in a compartment built from a single nucleotide, hydrolysis of both a small ester and the nucleic-acid backbone is suppressed. In this system, the reduced activity/availability of water in the dense phase appears to outweigh the concentrating effects that normally accelerate hydrolysis reactions. The stabilization of RNA we observe in nucleotide condensates could lead to new approaches to stabilize RNA therapeutics during distribution. The current generation of mRNA vaccines relies on lipid nanoparticle formulations whose shelf life is limited by RNA hydrolysis and requires ultra-cold storage [79]. A compartment built from the nucleotide monomers themselves, which stabilizes the enclosed RNA at ambient temperature, could offer an alternative route to room-temperature RNA formulations.

The most counterintuitive feature of the system studied here is that acidity, which is ordinarily destructive to nucleic acids, can actually preserve them by causing condensate formation. While recent comprehensive reviews on protein-free nucleic acid phase separation focus primarily on temperature-induced transitions and divalent cation coordination [29, 22], the potential of acidic environments remains overlooked due to conventional assumptions regarding acid-catalyzed degradation. This protonation-driven mechanism offers a remarkably straightforward and accessible pathway for nucleic acid compartmentalization, operating independently of high magnesium concentrations or thermal fluctuations. This partition establishes a sharp chemical selection mechanism: unprotected nucleic acids in the surrounding bulk solution are rapidly destroyed by acid-catalyzed hydrolysis, but the specific, water-depleted microenvironment inside the dense phase shields encapsulated polymers from degradation, even though the interior of nucleotide condensates is itself acidic, with a reported internal pH near 2.5 [50]. These observations may suggest that dense and acidic compartments in living cells could similarly host nucleic-acid chemistry that would be degradative in bulk solution.

The protection of nucleic acids from hydrolysis inside nucleotide condensates suggest that the same molecules that are able to encode genetic information in polymeric form are also capable of forming compartments that can protect such polymers. This could be a theoretical path to RNA stability in prebiotic settings which might also have produced RNA, placing the origin of biological compartments, biological information, and biological catalysis with prebiotically available molecules.

Although condensation or coacervation has long been considered a possible first compartment [14, 16, 17], it has not been clear what material these compartments would have been made from. Our results suggest that nucleotides can form compartments with biological potential. Such nucleotide compartments are particularly intriguing because nucleotides are the building blocks of the polymers carrying genetic information (RNA and DNA) in all cells today. It is also of interest that in extant cells, nucleotides serve multiple roles that have been essential since the last universal common ancestor [80]: not only are they the monomers for nucleic acid polymers, but also energy-carrying molecules and essential metabolic co-factors for metabolism. That these monomers which are so central to all organic life have the chemical properties needed to form a separate compartment implies the possibility of an even simpler prebiotic state, where a mixture of nucleotides formed compartments even before the emergence of polymeric RNA and DNA.

Theoretical models of RNA-condensate-catalyzed polymerization led to the idea that natural selection would emerge in such a system because diversity in condensate “phenotypes” would be encoded in the RNA “genotype” that determines the material properties of the condensate [48, 81]. Verstraeten et al. recently demonstrated this coupling experimentally in RNA droplets generated cotranscriptionally from a DNA template; mutations in the template produced distinct droplet phenotypes, including the emergence of long-lived vacuoles and droplet motility [82]. Although protection of the genetic information was not considered previously, protection of RNA polymers from hydrolysis is a condensate “phenotype” required in any such model for the origin of life. Furthermore, the idea that variation in RNA “genotypes” would lead to variation in condensate “phenotypes” is borne out in our observations of different material properties of, for example, ATP and GTP condensates. A “genotype” of mixed NTP condensates (e.g. G/A/C/U composition) is expected to result in a specific “phenotype” (e.g. condensate viscosity), *even in the absence of any RNA molecules longer than one nucleotide*. Thus, a population of nucleotide condensates with varying nucleotide composition is expected to show variation in material properties. If condensates can split and pass on their nucleotide composition, it seems theoretically possible that natural selection could begin in the absence of any polymer.

### Materials

Adenosine 5’-triphosphate (ATP) (A160197, Ambeed and FLUH99CD374F, Merck), Guanosine 5’-triphosphate (GTP) (G8877, Merck), Cytidine 5’-triphosphate (CTP) (C1506, Merck) Adenosine (A9251-1G, Merck), adenine hydrochloride (A17622.06-5G, Thermo Fisher Scientific), Adenosine 5’-diphosphate (ADP) (Merck, A2754) Sodium hexametaphosphate 96% (HMP) (305553-25G, Merck), Sodium triphosphate 85% (238502-25G, Merck), 1 M Magnesium chloride (MgCl_2_) (AM9530G-100 mL, Thermo Fisher Scientific) 1,4-butanediol, 99% (493732-1L, Merck), 1,3-propanediol 97% (201-02873-25 mL, FUJIFILM Wako), 1,2-propanediol 99% (164-04996, FUJIFILM Wako), 1,6-hexanediol (081-00435, FUJIFILM Wako), ethylene glycol (1.00949-1L, Merck), glycerol (G0912, Merck), Methanol (34860, Honeywell), ethanol (02870, Honeywell), p-Nitrophenyl phosphate (pNPP) (71768-5G, Merck), Methylene blue (133-06962, Wako), Resorufin (73144, Merck), Urea (different vendors, Merck, Thermo Fisher and Bio-Rad) Poly-L-lysine FITC Labeled 15-30 kDa (P3543, Merck), BioTracker ATP-Red Live Cell Dye (SCT045, Merck), *β*-BODIPY FL C_12_-HPC (D3792, Thermo Fisher Scientific), BODIPY 558/568 C12 (D3835, Thermo Fisher Scientific), UltraPure DNase/RNase-Free Distilled Water (10977035-500mL, Thermo Fisher), Tris-HCl 1 M pH 8.0 (AM9856-500mL, Thermo Fisher Scientific) and Tris-HCl 1 M pH 7.0 (AM9850G-500mL, Thermo Fisher Scientific), Formamide (F9037, Merck), RNA 2X gel loading buffer (R0641, Thermo Fisher Scientific), Acrylamide/N,N’-Methylenebisacrylamide 19:1, for biochemistry, 40% mix solution in water (330205000, Thermo Fisher Scientific). Oligonucleotides: FAM-ssRNA 12-mer, Cy3-ssDNA 16-mer, FAM-ssRNA 16-mer, FAM-ssRNA 20-mer, FAM-RNA hairpin 28-mer, DNA template for the hammerhead ribozyme and FAM-labeled RNA substrate were ordered from IDT (Integrated DNA Technologies, purified by desalting or by HPLC).

### Stock solutions

ATP was prepared at 500 mM or 1 M in ultrapure water (final pH was around 2.0); GTP and CTP at 200 mM; HMP at 200–500 mM; and sodium triphosphate at 200 mM. Adenine and Adenosine at 200 mM in acidic environment using HCl. MgCl_2_ was used from a 1 M stock (AM9530G-100 mL, Thermo Fisher Scientific). All nucleotide and phosphate stocks were prepared in UltraPure DNase/RNase-free water and stored at −20 *^◦^*C; once thawed, aliquots were used immediately and discarded (except for the oligonucleotides), and stocks were never refrozen or reused. All oligonucleotides (IDT) were resuspended to 100 µM in UltraPure DNase/RNase-free water.

### Condensate formation

Condensates were prepared in a total volume of 200 µL unless otherwise stated. Nucleotide condensates were formed by combining a nucleoside triphosphate (ATP, GTP or CTP) with MgCl_2_ and a short-chain diol or alcohol; phosphate condensates were formed with sodium hexametaphosphate (HMP) and sodium triphosphate. Standard conditions were 20-30 mM nucleotide or HMP, 20 mM MgCl_2_ and 50% (v/v) 1,4-butanediol. Acidic conditions (in the case of HMP partitioning test reported in Fig. 1) were established by adding 1 µL of 5 M HCl to the 200 µL sample (25 mM HCl), giving a pH below 3 (pH 2.2-2.5), measured after preparation with a microelectrode. The pH of the ATP condensates (without adding acid) prepared using 20 mM ATP, 20 mM MgCl_2_ and 50% 1,4 butanediol was 2.0-2.5.

### Diol and co-solvent survey

For the HMP diol comparison, HMP was used at 100 mM with ethylene glycol and at 20 mM with 1,2– and 1,4-diols, all at 50% (v/v); ethylene glycol at this concentration did not support HMP condensation at 20 mM. For the nucleotide co-solvent survey, ATP was combined with methanol (50-55%) or ethanol (40-45%) at 20 mM ATP, 20 mM MgCl_2_. The propanediol isomers were compared at 29 mM ATP, 30 mM MgCl_2_, with 60-65% 1,2-propanediol (aggregates) and 50% 1,3-propanediol (droplets). Adenine and adenosine were tested at 20-50 mM, ±MgCl_2_ 20 mM, in acid, with 50-70% 1,4-butanediol. Sodium triphosphate was tested at 20-60 mM with 20 mM MgCl_2_ and 50% 1,4– butanediol. GTP (10-20 mM, 25 mM MgCl_2_, 50-55% 1,4-butanediol) required acidification (10 µL of 0.5 M HCl in 200 µL) to form liquid droplets rather than aggregates. We note that GTP droplet formation proved technically challenging. The system has a strong tendency to form aggregates, and reagent purity likely plays a role. Allowing the samples to rest for a few hours also improved droplet formation. CTP: 20 mM, 20 mM MgCl_2_, 60-65% 1,4-butanediol.

### pH series using HMP

140 mM HMP, 50% ethylene glycol, 20 mM MgCl_2_, under acidic (125 mM HCl), neutral (100 mM Tris pH 7) or alkaline (125 mM NaOH) conditions.

### Microscopy and partitioning

Samples were imaged on an Olympus FluoView FV1200 confocal laser scanning microscope, on an inverted Zeiss Axio Observer.Z1/7 widefield fluorescence microscope, or on an EVOS M3000 imaging system (Thermo Fisher Scientific), in brightfield and fluorescence with 20× and 40× objectives. On the Zeiss system, images were acquired with an EC Plan-Neofluar20×/0.50 air objective, a FITC filter set (excitation 455–495 nm, emission 505–555 nm) and an AxioCam MR R3 camera (exposure time 170 ms, pixel size 0.3225 µm) using ZEN 2.3 (blue edition), and were analysed with ImageJ (v1.54t). A drop of sample (3–5 µL) was added to the glass slide and immediately visualized. Cargo was added before to form condensates under matched conditions and imaged after 30-120 min or after ON incubation. All partitioning assays were performed under acidic conditions to allow direct comparison of HMP and ATP (HCl was added only to HMP samples, as dissolving the acidic form of ATP in water naturally resulted in an acidic stock solution). Polymers and small organic molecules final concentrations: FAM-ssRNA 12-mer (5 µM), Cy3-ssDNA 16-mer (5 µM), methylene blue (0.01%), resorufin (50 µM), BioTracker ATP-Red Live Cell Dye (Sigma-Aldrich SCT045) 20 µM. Partitioning in Fig. 1a used 20 mM HMP or 20 mM ATP, 20 mM MgCl_2_, and 50% 1,4-butanediol. For the poly-L-lysine experiment in Fig. 1b, HMP was used with 1,4-butanediol and ATP with 35–40% ethanol. For amphiphiles, we used *β*-BODIPY FL C_12_-HPC 20 µM final concentration and BODIPY 558/568 C12 was used 25 µM. The images collected were analyzed using ImageJ. Brightness and contrast were adjusted for clear visualization of the molecules. Scale bars were 10 µm or 20 µm.

### Temperature cycling and reversibility

At neutral pH, ATP condensation was induced by cooling. Samples were prepared at room temperature with 32 mM ATP, 80 mM MgCl_2_, 100 mM NaCl, 125 mM Tris-HCl pH 8 and 30-35% 1,4-butanediol; because ATP is acidic, the final pH of the mixture was 6.7-7.2 even using high Tris-HCl concentration. Under these conditions no droplets form at room temperature. The resulting suspension (∼250 µL) was subjected to five cycles of 5 min on ice (∼1 *^◦^*C, with the tubes held in an aluminum block for microcentrifuge tubes that was kept on ice throughout to maintain the temperature) and 5 min at 30 *^◦^*C in the microplate reader. During each dissolution phase, 200 µL were transferred to a plate reader (MultiSkan SkyHigh microplate reader, Thermo Fisher Scientific) and OD_600_ recorded every 6 s for 5 min, then returned to the tube and placed back on ice. RNA partitioning used ssRNA-16mer at 5 µM.

### Hydrolysis assays

For the preparation of ATP condensates (20 mM ATP, 20 mM MgCl_2_, 50% 1,4-butanediol) and bulk controls (water + MgCl_2_; ATP + MgCl_2_; diol + ATP + MgCl_2_), components were added sequentially in the following order: ultrapure water, ATP (from a 1 M stock), MgCl_2_ (from a 1 M stock), pNPP (to a final concentration of 8 mM, from a 300 mM stock in water), and finally 1,4-butanediol (where applicable). Samples were briefly vortexed between additions to ensure homogeneity, and condensate formation was triggered by the addition of the diol in the condensate condition. The assay was performed at room temperature. For each time point, a separate 20 µL aliquot was prepared. At 0, 12, 24, and 48 h, the samples were diluted by adding 180 µL of 100 mM potassium phosphate buffer (pH 8.3) and mixed. The absorbance at 405 nm was measured using a Multiskan SkyHigh Microplate Reader (Thermo Fisher Scientific) at room temperature, protecting the samples from the light with aluminium foil. Data were analyzed using GraphPad Prism 11.1.0.

### Nucleic-acid fold prediction

Structures of the oligos were computed by the RNAfold WebServer using minimum free energy (MFE) and partition function with the avoid isolated base pairs option [75, 76]. RNA parameters (Turner model, 2004), DNA parameters (Matthews model, 2004) with 1 M salt at 37°C.

### Nucleic-acid protection

Single-stranded RNA (ssRNA) 12-mer, ssRNA 20-mer, and a 28-mer RNA hairpin were tested at a final concentration of 5 µM. Reagents were always added in the following order: water, ATP, MgCl_2_, RNA and, where required, diol. *Bulk (no diol) controls.* Samples contained 20 mM ATP and 20 mM MgCl_2_ and 5 µM RNA (final concentrations), without diol, and were incubated at 37 *^◦^*C. *ATP condensates.* Samples were prepared with 20 mM ATP, 20 mM MgCl_2_, 5 µM RNA and 50% diol (final concentrations). The diol was added last to trigger droplet formation, and samples were then incubated at 37 *^◦^*C after droplet formation. *Sub-threshold diol controls.* Samples contained 20 mM ATP, 20 mM MgCl_2_, 5 µM RNA and 35% (v/v) 1,4-butanediol (final concentrations), a concentration at which no condensates form, and were incubated at 37 *^◦^*C. To prevent evaporation in the 1.5 mL tubes, samples were monitored every 6–12 h and spun down when necessary (typically only for the bulk controls). Aliquots were collected over a period of 72 h or 10 days. The condensed phase was isolated by centrifugation, resuspended in 40 µL of water, and either stored at −40 *^◦^*C or immediately analyzed by denaturing polyacrylamide gel electrophoresis (PAGE). For the analysis, samples were diluted 1:5 in water, mixed with an equal volume of 2X loading buffer containing 50% formamide, heat-denatured, and cooled on ice. Finally, 2–4 µL of each sample was loaded onto the denaturing PAGE gel.

### Ribozyme assay

Reaction mixtures were assembled at 25 *^◦^*C by adding the components in the following order: water, ATP (condensate samples only), buffer and salts, the pre-folded ribozyme (carrying 5 mM MgCl_2_ and NaCl from the folding buffer) and the FAM-labeled substrate. MgCl_2_ was then added to bring its total concentration to 80 mM, which initiated the reaction (*t* = 0). For the condensate samples, 1,4-butanediol was added last, triggering droplet formation. Bulk samples were prepared in the same way, omitting ATP and diol.

Prior to the assay, the ribozyme was denatured and refolded in buffer (125 mM Tris-HCl pH 8.0, 100 mM NaCl, 2.5 mM KCl, 5 mM MgCl_2_) by heating at 90 *^◦^*C for 2 min and slow-cooling to 25 *^◦^*C at a rate of 0.1 *^◦^*C s*^−^*^1^ in a thermocycler. The FAM-labeled 26-mer substrate was used as supplied without a separate denaturation step, because the lack of secondary structure make a pre-folding step unnecessary. Reactions were initiated by combining the pre-folded ribozyme with the substrate in the final reaction mixture (either bulk buffer or ATP condensate), bringing MgCl_2_ to its final concentration of 80 mM.

Immediately after preparation, each 200 µL sample was divided into 40 µL aliquots. At each time point over 6 h (0, 30, 60, 120, 180, 240 and 360 min), condensate samples were centrifuged at 16,000×*g* for 1 min and the supernatant removed; the condensed phase was resuspended in UltraPure water and briefly heated in the presence of 50 mM EDTA to accelerate dissolution and arrest cleavage by chelating magnesium. 2× RNA loading dye (Thermo Fisher or prepared in-house) was then added and samples were denatured at 80 *^◦^*C for 2 min and placed immediately on ice. Products were resolved on denaturing urea-PAGE. The fraction cleaved was quantified as the ratio of cleaved product to total RNA (cleaved product + substrate), and substrate recovery as the decrease in fraction cleaved from its peak to the endpoint. *n* = 3 independent experiments per condition.

### Denaturing PAGE

Denaturing urea–PAGE gels (20% acrylamide, 19:1, ∼8 M urea) were prepared per 8 mL with 3.8 g urea, 800 µL 10× TBE, made up to 8 mL with 40% acrylamide/bis, and polymerized with 40 µL 10% APS and 10 µL TEMED. Samples were mixed with loading buffer containing 50% formamide and denatured before loading. RNA samples were heated at 70–80 *^◦^*C for 2 min. Samples were cooled before loading, and 2–4 µL were loaded per lane. Gels were run on a Bio-Rad electrophoresis system and imaged on an iBright CL-1500 imaging system (Invitrogen, Thermo Fisher Scientific). Band intensities were quantified for the main band and all degradation products in the same lane using ImageJ. The product of area measured and mean intensity was used to calculate the intact fractions of the main band after subtraction of background.

### Statistical analysis

The pNPP hydrolysis data (Fig. 2f,g) were analyzed by Welch’s one-way ANOVA followed by Dunnett’s T3 multiple-comparisons test, which does not assume equal variances across groups (GraphPad Prism 11.1.0). Sample sizes: Water + MgCl_2_, *n* = 13; ATP + MgCl_2_ (bulk), *n* = 12; ATP condensates, *n* = 13; diol + ATP + MgCl_2_ (bulk, sub-threshold), *n* = 8. At 24 h, Welch’s *W* (3, 20.83) = 86.11, *p <* 0.0001. The condensate differed from every bulk condition: versus Water + MgCl_2_, *t*(13.74) = 10.21, adjusted *p <* 0.0001; versus ATP + MgCl_2_, *t*(12.74) = 9.08, adjusted *p <* 0.0001; versus diol + ATP + MgCl_2_ (sub-threshold), *t*(13.97) = 12.59, adjusted *p <* 0.0001. At 48 h, Welch’s *W* (3, 18.75) = 290.3, *p <* 0.0001. The condensate differed from every bulk condition: versus Water + MgCl_2_, *t*(13.38) = 18.78, adjusted *p <* 0.0001; versus ATP + MgCl_2_, *t*(11.73) = 11.84, adjusted *p <* 0.0001; versus diol + ATP + MgCl_2_ (sub-threshold), *t*(10.07) = 23.87, adjusted *p <* 0.0001. Differences among bulk conditions were small and partly non-significant at both time points.

### Molecular dynamics simulation

To study ATP and DNA phase separation, we performed molecular dynamics simulation using a coarse-rained description of ATP and DNA, similar to what has been done for RNA and poly(ADP-ribose) (PAR) [73, 83, 84, 85]. ATP is modeled using a five bead representation, where each bead is located at the center-of-mass of its corresponding group: one for each phosphate group, one for sugar and another for base. The energy function for ATP is: *E*_ATP_ = *E*_bond_ + *E*_angle_ + *E*_DH_ + *E*_ev_. *E*_bond_ and *E*_angle_ are the bonded and angle interactions which are modeled using harmonic potentials. *E*_DH_ is the electrostatic interaction between two phosphate beads which is modeled using Debye-Hückel screening potential. *E*_ev_ is the excluded volume interaction between two non-bonded beads which is modeled using the Weeks-Chandler-Anderson (WCA) potential. Because ATP contains the same functional groups as the monomer unit of PAR, we used the same parameter set recently developed for PAR [85, 86] for ATP.

DNA is modeled using a three bead coarse-grained description for phosphate, sugar and base [84]. In our simulations, Mg^2+^ ions are explicitly present within the simulation box. The interaction between phosphate groups and Mg^2+^ ions are calculated as described elsewhere [73], using *E*_P-Mg_ = *W* (*r*) + [*E*_DH_ − *W* (*r*)] exp(−*a*^2^*/r*^2^) where *W* (*r*) is the potential of mean force obtained from RISM theory [87, 88]. This potential can distinguish between direct and solvent mediated binding modes of ions which depends on temperature [89], ion concentration [85, 86, 90, 91] and nucleic acid structure [92]. We performed Langevin dynamics simulations using a time step of 2 fs and snapshots are saved after every 20 ps. The initial structure for simulation is generated by randomly inserting 60 ssDNA, 1000 ATP and 2200 Mg^2+^ ions within a cubic box of length 700 Å. All simulations are performed using OpenMM [93].

To estimate the number of water molecules in the dilute and dense phases, we calculated the solvent-accessible surface area (SASA) considering only phosphate beads in the respective phase using the freeSASA Python package [94]. Specifically, a spherical probe with a radius of r = 1.4 Å, approximately the radius of a water molecule, was rolled over the surface of each phosphate bead individually to calculate its SASA, *S_p_*. The average number of water molecules associated with each phosphate is then approximated as *N*_water_ = ⟨*S_p_*⟩ / *πr*^2^.

## Author contributions

S.H. and M.D. conceived the project. S.H., R.N. and M.D. performed experiments. L.B. and H.T.N performed molecular simulations. All the authors contributed to the manuscript. M.D. supervised the project with inputs from all the authors.

## Acknowledgements

We thank Rohit Pappu, Jerelle Joseph, and Andrei Lupas for their comments on the manuscript. We thank Walter Gilbert for inspiring discussions. M.D. gratefully acknowledges the University of Perugia for the support. H.T.N. gratefully acknowledges financial support from the University at Buffalo. R.N. thanks the Foundation of Kinoshita Memorial Enterprise (FY2026) for financial support. Computational resources were provided by the Center for Computational Research at the University at Buffalo.

