## Supplementary Information for "Unexpected properties of nucleotide condensates could facilitate the RNA world"

### Supporting Information Text

**Table S1.** Oligonucleotides used in this study.

| Name | Sequence (5′ to 3′) | Label | Supplier | Stock |
|---|---|---|---|---|
| ssRNA 12-mer | GUGUCGCUUUCG | 5′-FAM | IDT | 100 µM |
| ssDNA 16-mer | TTAAGTGTCGCTTTCG | 5′-FAM | IDT | 100 µM |
| ssDNA 16-mer | TTAAGTGTCGCTTTCG | 5′-Cy3 | IDT | 100 µM |
| ssRNA 16-mer | UAUGCGCAUGCCUUCG | 5′-FAM | IDT | 100 µM |
| ssRNA 20-mer | AAAAUGAACUGAACUUGUUU | 5′-FAM | IDT | 100 µM |
| RNA hairpin 28-mer | UUAAUUGGCCGCAUGCCUUCGGGCAUGC | 5′-FAM | IDT | 100 µM |
| RNA hairpin 23-mer | UAUGCGCAUGCCUUCGGGCAUGC | 5′-FAM | IDT | 100 µM |
| Hammerhead template, fwd | TAATACGACTCACTATAGGGCGAGGTACATCCAGCTGACGAGTCCCAAATAGGACGAAATGCC | unlabeled | IDT | 100 µM |
| Hammerhead template, rev | GGCATTTCGTCCTATTTGGGACTCGTCAGCTGGATGTACCTCGCCCTATAGTGAGTCGTATTA | unlabeled | IDT | 100 µM |
| Hammerhead substrate | AAAAGGCAUCCUGGAUUCCACUCGCC | 5′-FAM | IDT | 100 µM |
| Hammerhead substrate, poly-A tailed | AAAAAAAAAAAAAGGCAUCCUGGAUUCCACUCGCC | 5′-FAM | IDT | 100 µM |

All oligonucleotides were synthesized by IDT and resuspended to 100 µM. The fwd and rev strands anneal to a double-stranded T7 template; the catalytic hammerhead RNA is produced by *in vitro* transcription. Structures were computed by the RNAfold WebServer using minimum free energy (MFE) and partition function with the avoid isolated base pairs option. RNA parameters (Turner model, 2004), DNA parameters (Matthews model, 2004) with 1 M salt at 37°C.

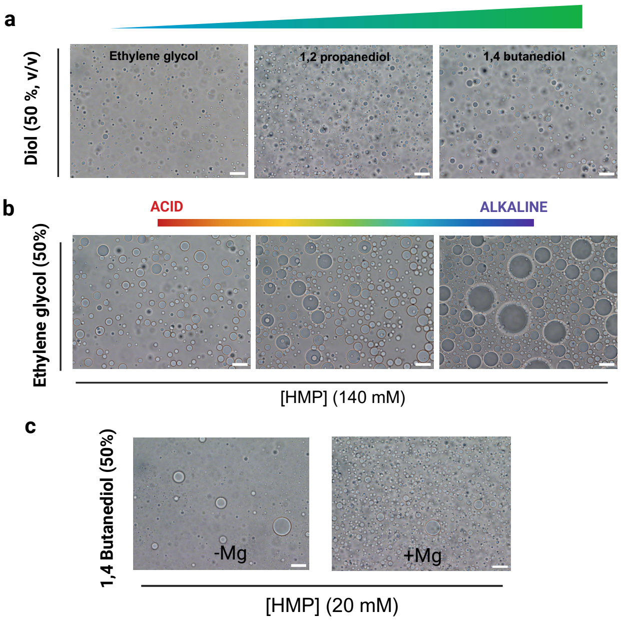


**Fig. S1. Phosphate (HMP) phase separation is robust to conditions.** **(a)** Diol survey. HMP forms droplets with a range of short-chain diols (ethylene glycol, 1,2-propanediol, 1,3-propanediol, 1,4-butanediol and 1,6-hexanediol) at 50% (v/v), in the presence of 20 mM $MgCl_2$. HMP was used at 100 mM with ethylene glycol and at 20 mM with the other diols. **(b)** pH range. HMP droplets (140 mM HMP, 50% ethylene glycol, 20 mM $MgCl_2$) form across acidic (125 mM HCl), neutral (100 mM Tris-HCl, pH 7) and basic (125 mM NaOH) conditions. **(c)** Magnesium dependence. HMP droplets (20 mM HMP, 50% 1,4-butanediol) form both in the presence of 20 mM $MgCl_2$ and in its absence. Scale bars, 20 µm.

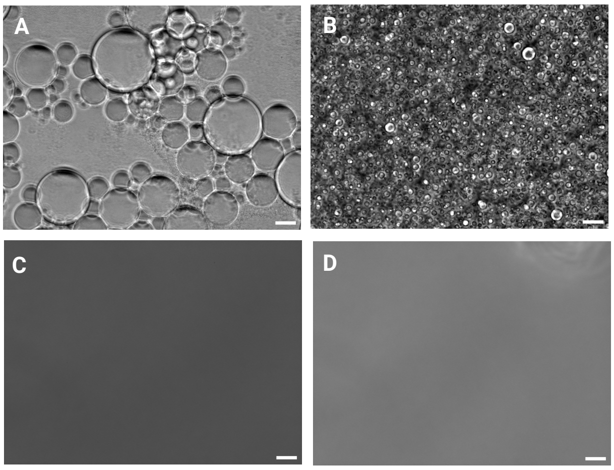


**Fig. S2. Phase separation requires the phosphate, not the base.** Brightfield images of candidate components tested under standard condensation conditions. **(a)** Sodium triphosphate (40 mM) in 50% 1,4-butanediol, without $MgCl_2$. **(b)** Sodium triphosphate (40 mM) in 50% 1,4-butanediol with 20 mM $MgCl_2$. **(c)** Adenine (40 mM) with 20 mM $MgCl_2$ and 50% 1,4-butanediol. **(d)** Adenosine (40 mM) with 20 mM $MgCl_2$ and 50% 1,4-butanediol. Scale bars, 20 µm.

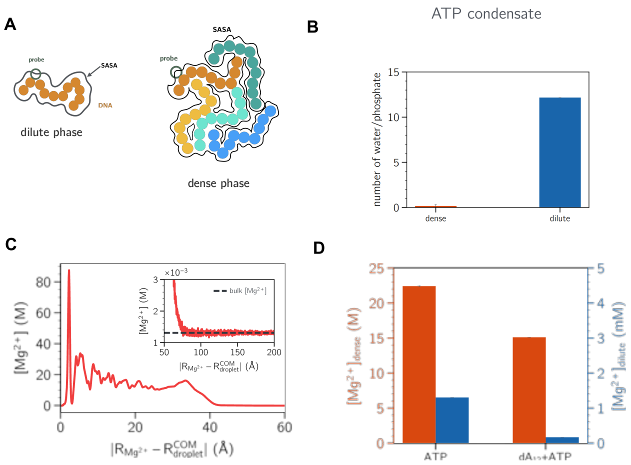


**Fig. S3. Water content and magnesium distribution within the condensate (simulations). (a)** Schematic of the solvent-accessible surface area (SASA) approach used to estimate the number of water molecules bound per phosphate for dilute and dense phase. The DNA chains are represented in different colors. **(b)** Estimated bound water per phosphate in the dilute and dense phases; water bound per phosphate is markedly lower in the dense phase. **(c)** Radial $Mg^{2+}$ concentration as a function of distance from the condensate centre of mass, decaying to the bulk value (dashed line) in the inset. **(d)** $Mg^{2+}$ concentration in the dense phase decreases upon adding ssDNA to the ATP condensate. Concentrations from coarse-grained simulations are relative rather than absolute; DNA is modelled as homopolymers ($dA_8$, $dA_{12}$).

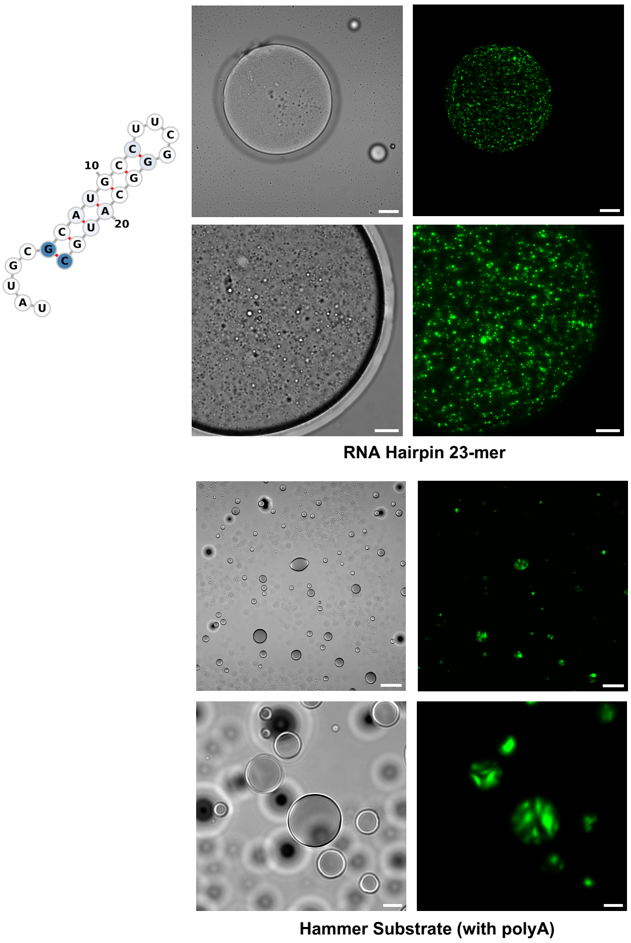


**Fig. S4. Structured and tailed RNAs organize non-uniformly within ATP condensates.** Confocal fluorescence of FAM-labelled RNAs concentrated inside ATP condensates (20 mM ATP, 20 mM $MgCl_2$, 50% 1,4-butanediol, acidic pH). **(a)** RNA hairpin 23-mer (`UAUGCGCAUGCCUUCGGGCAUGC`), concentrating into discrete intradroplet puncta distributed throughout the droplet volume. **(b)** Hammerhead substrate carrying a 5′ poly-A tail ($A_{13}$`GGCAUCCUGGAUUCCACUCGCC`, 35-mer; forming a radial/striated internal pattern. Together with the hairpin 28-mer shown in Fig. 4, these images indicate that structured and tailed RNAs do not fill the droplet uniformly but organize into sub-droplet domains. RNA at 5 μM. Scale bars, 20 μm.

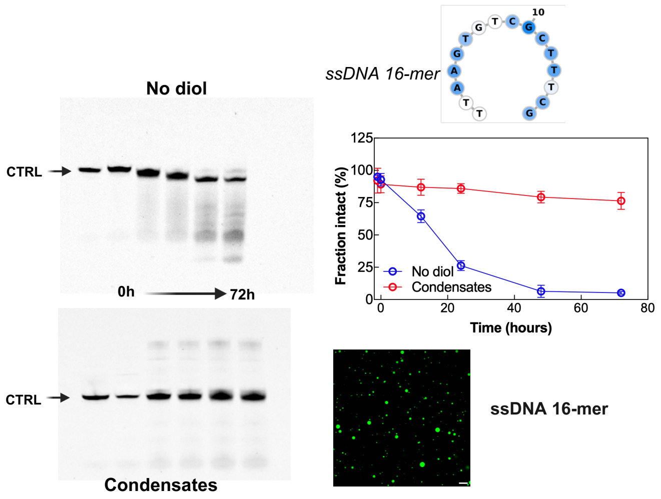


**Fig. S5. Single-stranded DNA is also protected from acid hydrolysis inside ATP condensates.** Left, negative images of fluorescence signals after denaturing urea-PAGE for the FAM-labelled ssDNA 16-mer in the no-diol control (water + ATP + $MgCl_2$, acidic pH; top) and inside ATP condensates (20 mM ATP, 20 mM $MgCl_2$, 50% 1,4-butanediol, acidic pH; bottom), over 72 h at 37 °C, with untreated control (CTRL). Top right, secondary-structure prediction of the ssDNA 16-mer (RNAfold, DNA Mathews 2004 parameters, 1 M salt, 37 °C). Middle right, fraction of intact ssDNA 16-mer over 72 h, no-diol control (blue) versus condensate (red); mean ± SD, $n = 3$. In the no-diol control the ssDNA 16-mer drops to ∼39% by 72 h, whereas inside the condensate it remains above 94%. Bottom right, confocal fluorescence showing the FAM-labelled ssDNA 16-mer concentrated into ATP condensates. DNA lacks the 2'-OH that drives the intramolecular transesterification mechanism of RNA cleavage, so its acid-driven degradation follows a different chemistry; the condensate stabilises both nucleic acids to a comparable extent, consistent with a protection mechanism intrinsic to the dense-phase environment. Oligonucleotide at 5 $\mu$M; acidic conditions (pH 2–2.3). Scale bar, 20 $\mu$m.

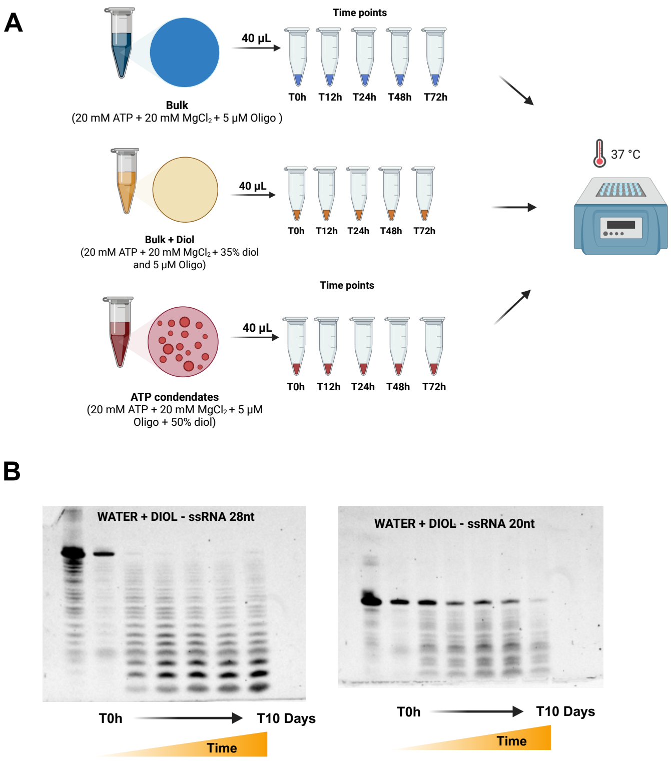


**Fig. S6. The diol does not protect nucleic acids in the absence of condensation. (a)**, Schema of the experimental set-up. **(b)** Full-length fraction of the ssRNA 20-mer and RNA hairpin 28-mer over 10 days at 37 °C in bulk with diol (water + ATP + $MgCl_2$ + 35% 1,4-butanediol, no condensation).

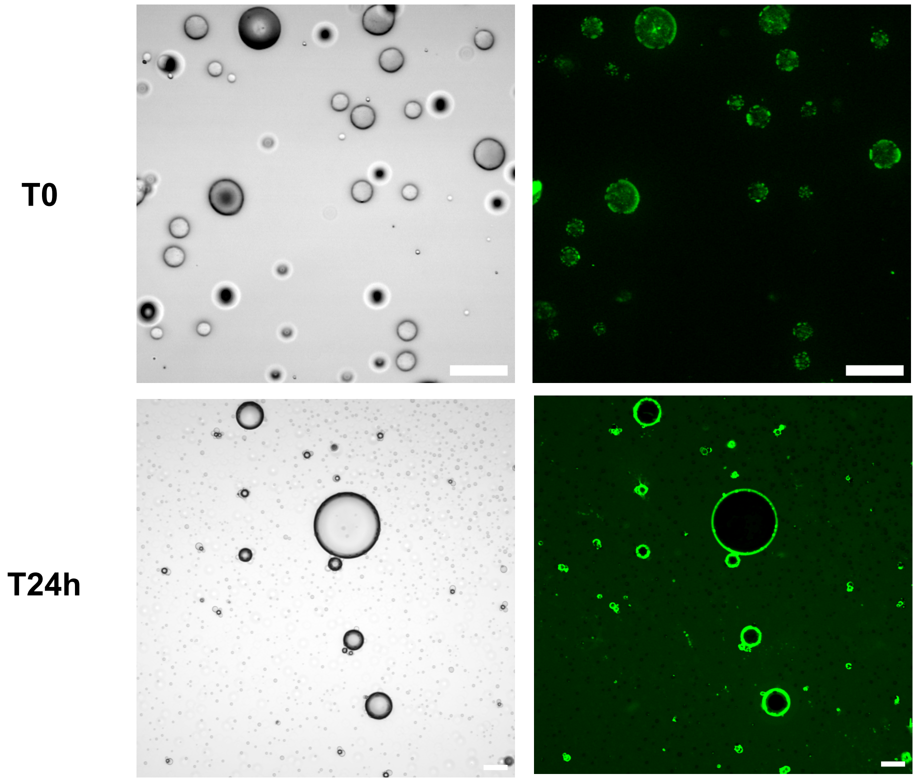


**Fig. S7. The phospholipid redistributes from internal foci to a surface layer over hours.** Confocal and brightfield imaging of ATP condensates (40 mM ATP, 20 mM $MgCl_2$, 50% 1,4-butanediol, acidic pH) incubated with the phospholipid $\beta$-BODIPY FL $C_{12}$-HPC (20 µM). **(a)** At T0, the phospholipid is distributed inside the droplet as internal puncta, with little signal at the interface. **(b)** By 24 h, the phospholipid has redistributed to a bright ring at the droplet surface, leaving the interior dark. Scale bars, 20 µm.
